# BMP4 restricts axonal ramification and bouton density in cholinergic projections from the basal forebrain

**DOI:** 10.64898/2026.09.30.755469

**Authors:** Takahito Higashi, Yasushi Kobayashi, Akiko Seki, Shigeo Okabe, Kiyomasa Nishii

## Abstract

Axonal projections and precise synapse formation are essential for the establishment of functional neural circuits, and their dysregulation contributes to various psychiatric and neurodegenerative disorders. Although numerous extracellular cues secreted from surrounding tissues regulate axon guidance and synapse formation, how neurons locally regulate terminal patterning within their projection fields remains poorly understood. Bone morphogenetic protein 4 (BMP4) is a classical morphogen involved in early neural tube patterning, but its postnatal function remains unclear. A previous *in vitro* study using hippocampal glutamatergic neurons demonstrated that BMP4 can act locally on axons to suppress synapse formation. We therefore hypothesized that BMP4 restricts axonal ramification and terminal formation after axons reach their targets.

Here, we tested this hypothesis in basal forebrain cholinergic neurons, which project extensively throughout the neocortex and are essential for higher brain functions, including arousal and attention. We generated cholinergic neuron-specific *Bmp4* conditional knockout (*Bmp4* cKO) mice and performed adult analyses in males. In *Bmp4* cKO mice, cholinergic innervation appeared denser in the neocortex, hippocampus, and amygdala, and quantification in the somatosensory cortex confirmed an increased area fraction of cholinergic terminals in most layers. In contrast, the number and rostrocaudal distribution of basal forebrain cholinergic neurons were unchanged. Reconstruction of virally labeled cholinergic axons in the neocortex revealed more branch points, most prominently in layer V, and an overall increase in the density of axonal varicosities. These findings demonstrate that BMP4 restricts axonal ramification and bouton density in basal forebrain cholinergic projections.

**Significance Statement:** Cholinergic projections from the basal forebrain are critical for cortical function, yet the mechanisms that regulate their axonal ramification and bouton density remain poorly understood. Using cholinergic neuron-specific *Bmp4* conditional knockout mice, we show that *Bmp4* deletion increases cholinergic axonal ramification and bouton density in the neocortex without altering basal forebrain cholinergic neuron number. These findings identify BMP4 as a negative regulator of cholinergic innervation.

## Introduction

During development, neurons extend long-range axons to connect with specific target populations (Sanes and Yamagata, 1999). Within their target regions, these axons branch extensively and form synapses with appropriate partners (Gibson and Ma, 2011). The number and distribution of these synapses must be precisely controlled to ensure proper neural function (Heckman and Doe, 2021), and disrupted synaptic integrity is associated with psychiatric and neurological disorders, including schizophrenia, autism, and dementia (Penzes et al., 2011). Although numerous extracellular cues secreted from surrounding tissues regulate axon guidance and synapse formation (Chédotal, 2019; Südhof, 2021), the mechanisms by which neurons locally regulate axonal ramification and synaptic connectivity within their target regions remain poorly understood. In particular, little is known about neuron-derived signals that act after axons reach their target regions.

Bone morphogenetic protein 4 (BMP4), a member of the transforming growth factor-β (TGF-β) superfamily, is a classical morphogen involved in early neural tube patterning and other developmental processes, including cell fate specification and morphogenesis (Miyazono et al., 2010; Bond et al., 2012; Katagiri and Watabe, 2016). BMP4 binds to type I (BMPRIA/ALK3 and BMPRIB/ALK6) and type II (BMPRII) receptors, resulting in Smad phosphorylation and downstream transcriptional regulation (Rosenzweig et al., 1995; Miyazono et al., 2010; Katagiri and Watabe, 2016). Beyond early patterning, BMP and TGF-β signaling through Smad proteins also contributes to the formation of specific neural circuits, such as connections between the midbrain and striatum (Terauchi et al., 2023), auditory brainstem circuits (Xiao et al., 2013), and interneurons in the dorsal spinal cord (Andrews et al., 2017). BMP4 remains broadly expressed in the postnatal central nervous system (Mikawa et al., 2006; Higashi et al., 2018), and a previous *in vitro* study using hippocampal glutamatergic neurons demonstrated that BMP4 is secreted locally from axon terminals and suppresses synapse formation (Higashi et al., 2018). However, whether BMP4 regulates axonal ramification and bouton density *in vivo* remains unknown.

Basal forebrain cholinergic neurons provide a suitable system to address this question. These neurons regulate attention, learning, memory, and arousal (Wenk, 1997; Hasselmo, 1999; Xu et al., 2015; Guo et al., 2019). They are conventionally classified into the Ch1–Ch4 cell groups, located in the medial septum (Ch1), the vertical (Ch2) and horizontal (Ch3) limbs of the diagonal band of Broca, and the nucleus basalis of Meynert/substantia innominata (Ch4), and collectively provide the major cholinergic projections to the neocortex, hippocampus, and amygdala (Mesulam et al., 1983; Woolf, 1991; Wenk, 1997; Mechawar and Descarries, 2001; Janiesch et al., 2011; Ballinger et al., 2016; Li et al., 2018; Takács et al., 2018). Most cholinergic neurons in the ventral pallidum resemble the burst-firing type of basal forebrain cholinergic neurons and project to the medial prefrontal cortex and basolateral amygdala (Schlingloff et al., 2026). In their target regions, cholinergic axons form highly branched arbors bearing numerous varicosities (Mechawar and Descarries, 2001), which makes them well suited for examining how axonal ramification and bouton density are regulated. Moreover, whereas intrinsic cholinergic interneurons are concentrated in regions such as the striatum, nucleus accumbens, olfactory tubercle, and islands of Calleja (Kandel et al., 2000), cholinergic terminals in the neocortex, hippocampus, and amygdala derive predominantly from basal forebrain projection neurons (Mesulam et al., 1983; Ballinger et al., 2016). In this study, we generated cholinergic neuron-specific *Bmp4* conditional knockout (*Bmp4* cKO) mice using choline acetyltransferase (ChAT)-Cre mice (Gong et al., 2007) and investigated whether BMP4 regulates cholinergic axonal ramification and bouton density.

## Materials and Methods

### Ethics Statement

All animal procedures were approved by the Animal Experiment Ethics Committee of the National Defense Medical College and were conducted in accordance with institutional guidelines and the Guidelines for Proper Conduct of Animal Experiments (Science Council of Japan, 2006). All recombinant DNA experiments were approved by the Recombinant DNA Experiment Ethics Committee of the National Defense Medical College and were conducted in accordance with institutional biosafety regulations.

### Animals

Female ChAT-Cre transgenic mice (Tg(Chat-cre)GM24Gsat; MMRRC:017269-UCD; Gong et al., 2007) were first crossed with male *Bmp4* floxed mice (*Bmp4^floxneo/floxneo^*; hereafter *Bmp4^fl/fl^*; The Jackson Laboratory, IMSR_JAX:016878; Liu et al., 2004) to generate *Bmp4^fl/+^*; ChAT-Cre offspring. Female *Bmp4^fl/+^*; ChAT-Cre mice were then crossed with male *Bmp4^fl/fl^* mice to generate *Bmp4^fl/fl^*; ChAT-Cre mice. Finally, female *Bmp4^fl/fl^*; ChAT-Cre mice were mated with male *Bmp4^fl/fl^* mice to generate experimental littermates.

Male *Bmp4^fl/fl^* littermates lacking the ChAT-Cre transgene served as controls, whereas male *Bmp4^fl/fl^*; ChAT-Cre littermates served as *Bmp4* cKO mice. Unless otherwise stated, male littermates aged 3–4 months were used for all experiments. For the analysis of Smad1/5 phosphorylation (Fig. 2D–F), mice of either sex at postnatal days 4–5 were used.

For the Cre-dependent viral labeling experiments shown in Fig. 6, *Bmp4^+/+^*; ChAT-Cre mice were used as controls because Cre recombinase is required for expression of the enhanced green fluorescent protein (EGFP) transgene carried by the AAV9-pCAG-FLEX-EGFP-WPRE vector (Addgene, 51502-AAV9, USA; Oh et al., 2014). *Bmp4^+/+^*; ChAT-Cre mice were obtained by crossing the *Bmp4^fl/+^*; ChAT-Cre mice described above with C57BL/6J mice (CLEA Japan, Japan).

To verify Cre recombinase activity in cholinergic neurons, ChAT-Cre mice were crossed with mT/mG reporter mice (*Gt(ROSA)26Sor^tm4(ACTB-tdTomato,-EGFP)Luo^*/J; IMSR_JAX:007576; Muzumdar et al., 2007).

Mice were group-housed under standard laboratory conditions with a 12 h light/dark cycle (lights on at 07:00, defined as Zeitgeber time [ZT] 0; lights off at 19:00) at 24 ± 1°C and had *ad libitum* access to food and water. Perfusions and tissue collections were performed during the light phase (ZT0–ZT12), predominantly in the afternoon.

### Tissue Preparation and Immunolabeling

Adult male mice aged 3–4 months were anesthetized with a mixture of medetomidine (0.75 mg/kg; Zenoaq, Japan), midazolam (4.0 mg/kg; Sandoz, Japan), and butorphanol (5.0 mg/kg; Meiji, Japan) (modified from Kawai et al., 2011) and transcardially perfused with 4% paraformaldehyde (PFA) in phosphate-buffered saline (PBS, pH 7.4). The brains were removed and post-fixed for 24 h at 4°C. For Smad1/5 phosphorylation analysis (Fig. 2D–F), mice of either sex at postnatal days 4–5 were anesthetized by hypothermia on ice and decapitated, and the brains were immediately removed and immersion-fixed in 4% PFA in PBS for 24 h at 4°C. All brains were then cryoprotected overnight in 20% glycerol dissolved in 0.1 M phosphate buffer (PB, pH 7.4).

Coronal sections were cut at 50 μm using a sliding microtome (HM440E, Carl Zeiss, Germany) for free-floating immunostaining or at 5–7 μm using a cryostat (CryoStar NX70, Epredia, USA) for immunofluorescence staining. For sliding microtome sectioning, brains were frozen on dry ice. For cryostat sectioning, brains were embedded in Cryomount (Muto Pure Chemicals, Japan) and frozen in isopentane precooled in a dry ice–ethanol bath. Brains from mice at postnatal days 4–5 were sectioned using the cryostat. The primary and secondary antibodies used in this study are listed in Tables 1 and 2, respectively.

**Table 1.**
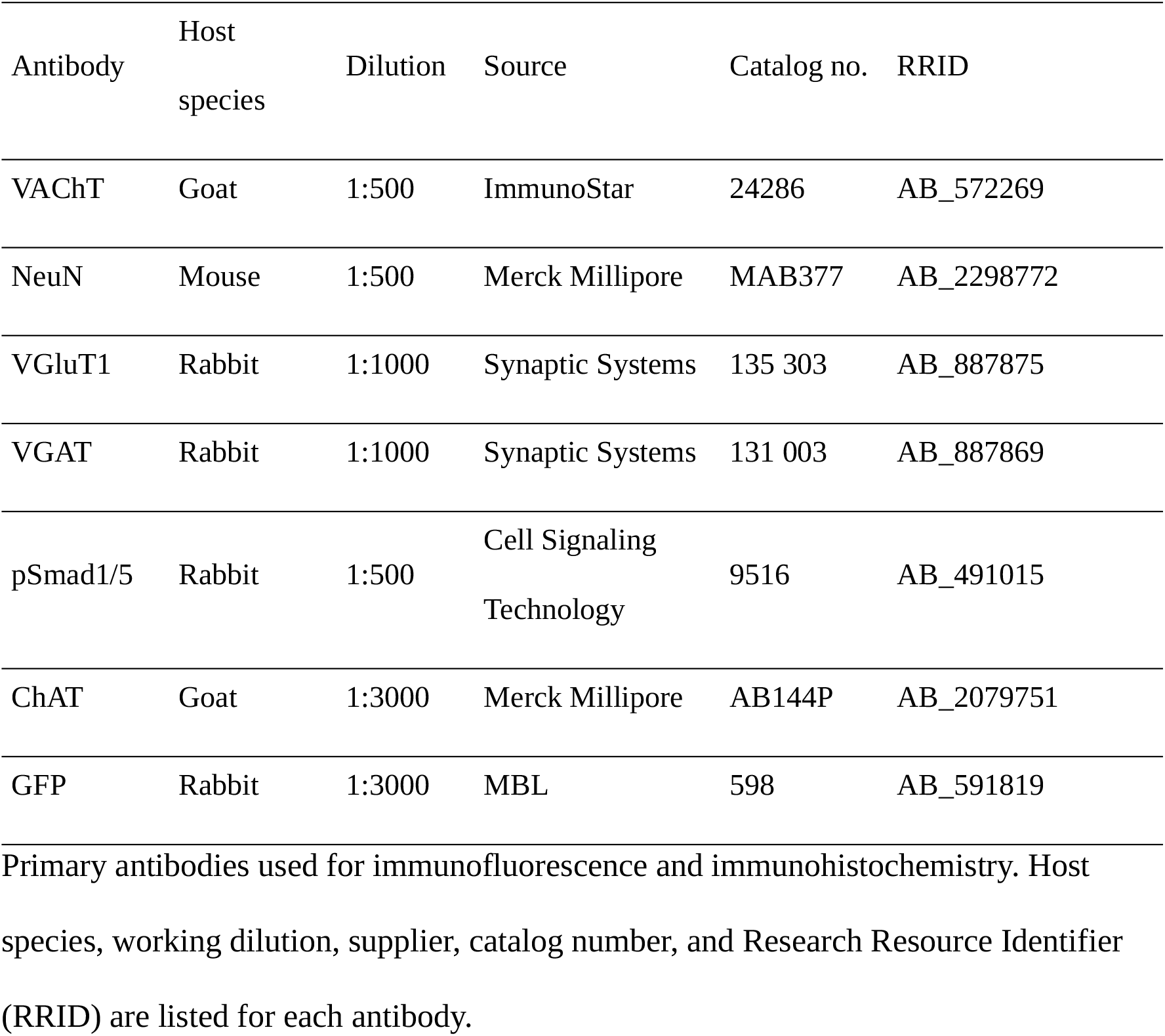
Primary antibodies used in this study.

| Antibody | Host species | Dilution | Source | Catalog no. | RRID |
|---|---|---|---|---|---|
| VAChT | Goat | 1:500 | ImmunoStar | 24286 | AB_572269 |
| NeuN | Mouse | 1:500 | Merck Millipore | MAB377 | AB_2298772 |
| VGluT1 | Rabbit | 1:1000 | Synaptic Systems | 135 303 | AB_887875 |
| VGAT | Rabbit | 1:1000 | Synaptic Systems | 131 003 | AB_887869 |
| pSmad1/5 | Rabbit | 1:500 | Cell Signaling Technology | 9516 | AB_491015 |
| ChAT | Goat | 1:3000 | Merck Millipore | AB144P | AB_2079751 |
| GFP | Rabbit | 1:3000 | MBL | 598 | AB_591819 |

**Table 2.** Secondary antibodies used in this study.

| Antibody | Conjugate | Dilution | Source | Catalog no. | RRID |
|---|---|---|---|---|---|
| Donkey anti-goat IgG | Alexa Fluor 488 | 1:500 | Thermo Fisher Scientific | A-11055 | AB_2534102 |
| Donkey anti-goat IgG | Alexa Fluor 546 | 1:500 | Thermo Fisher Scientific | A-11056 | AB_2534103 |
| Donkey anti-mouse IgG | Alexa Fluor 546 | 1:500 | Thermo Fisher Scientific | A-10036 | AB_2534012 |
| Donkey anti-rabbit IgG | Alexa Fluor 488 | 1:500 | Thermo Fisher Scientific | A-21206 | AB_2535792 |
| Donkey anti-goat IgG | Cy5 | 1:500 | Jackson ImmunoResearch | 705-175-147 | AB_2340415 |
| Rabbit anti-goat IgG | Biotin | 1:3000 | Vector Laboratories | BA-5000 | AB_2336126 |
| Goat anti-rabbit IgG | Biotin | 1:3000 | Vector Laboratories | BA-1000 | AB_2313606 |

Fluorophore-conjugated antibodies (Alexa Fluor 488, Alexa Fluor 546, and Cy5) were used for immunofluorescence, and biotinylated antibodies were used for immunohistochemistry with the avidin–biotin complex method. Conjugate, working dilution, supplier, catalog number, and Research Resource Identifier (RRID) are listed for each antibody.

Cryostat sections for immunofluorescence staining were subjected to antigen retrieval in citrate buffer (pH 6.0) at 95°C for 1 h. For the viral labeling experiments (Fig. 6), free-floating 50-μm sections were processed without antigen retrieval. Sections were then blocked with 5% normal horse serum in PBS containing 0.5% Triton X-100 at room temperature. Cryostat sections were incubated overnight at 4°C with the appropriate primary antibodies (Table 1), whereas sections used for viral labeling were incubated for 48 h at 4°C with anti-GFP and anti-ChAT antibodies (Table 1). After three washes in PBS, sections were incubated with the appropriate fluorophore-conjugated secondary antibodies (Table 2). For the viral labeling experiments, secondary antibodies were applied for 24 h at 4°C.

For immunohistochemistry, free-floating 50-μm sections were incubated with the appropriate primary antibodies (Table 1) for 48 h at 4°C. After three washes in PBS, sections were incubated with the appropriate biotinylated secondary antibodies (Table 2) for 24 h at 4°C, followed by incubation with the VECTASTAIN Elite ABC reagent (Vector Laboratories, USA) overnight at 4°C. Immunoreactivity was visualized using a nickel-enhanced 3,3′-diaminobenzidine (DAB) solution containing 0.002% H₂O₂, 0.03% DAB (Dojindo, Japan), and 0.03% nickel chloride in Tris-buffered saline.

### Stereotaxic Viral Injection

Male mice aged 3–4 months were placed in a stereotaxic apparatus (Narishige, Japan) for intracranial injection of adeno-associated virus serotype 9 (AAV9)-pCAG-FLEX-EGFP-WPRE (Addgene, 51502-AAV9, USA). The viral vector (1.65 × 10¹Z vg/mL) was injected into two basal forebrain sites at different rostrocaudal levels to achieve broad labeling. A glass micropipette (World Precision Instruments, USA) attached to a 5-μL gastight microsyringe with a glass Luer tip (Series H; ILS Innovative Laborsysteme, Germany) was guided to each site using coordinates based on a mouse brain atlas (Paxinos and Franklin, 2001): AP −0.1 mm, ML +1.3 mm, and DV −5.5 mm for the rostral site and AP −0.7 mm, ML +1.8 mm, and DV −3.5 mm for the caudal site (AP and ML relative to bregma; DV relative to the dura). An injection volume of 100 nL per site and an injection speed of 5 nL/s were controlled using a programmable Micro4 controller (World Precision Instruments, USA). The micropipette remained *in situ* for 5 min after each injection. Mice were transcardially perfused with 4% PFA 5 days after viral injection.

### Microscopy and Image Analysis

Fluorescence images were acquired using an Olympus IX83 inverted microscope (Olympus/Evident, Japan), a Nikon Eclipse 80i microscope (Nikon Solutions, Japan), and a Leica SP8 confocal laser-scanning microscope (Leica Microsystems, Germany). Image processing and quantitative analyses were performed using Fiji (ImageJ, NIH, USA).

For fluorescence quantification, background fluorescence was measured in regions within each image that showed no visible specific fluorescence and appeared dark on visual inspection. The threshold was defined separately for each image as the mean background intensity plus five standard deviations. Pixels with intensities above this threshold were considered positive, and background-subtracted fluorescence intensity was calculated by subtracting the threshold value from the intensity of each suprathreshold pixel. For vesicular glutamate transporter 1 (VGluT1) and vesicular GABA transporter (VGAT) analyses, the area fraction was calculated as the area above the fluorescence threshold divided by the total area analyzed.

For phosphorylated Smad1/5 (pSmad1/5) analysis, images were acquired from the basal forebrain of both hemispheres. The fluorescence intensity of pSmad1/5 was measured in individual vesicular acetylcholine transporter (VAChT)-positive and VAChT-negative neurons, and the mean fluorescence intensity of VAChT-negative neurons was calculated separately for each image. The fluorescence intensity of each neuron was then expressed relative to the mean fluorescence intensity of VAChT-negative neurons within the same image. Regions in which VAChT-positive neurons could not be identified were excluded from the analysis.

For quantification of cholinergic terminals, sections immunostained with an anti-VAChT antibody were analyzed. The VAChT-positive area fraction was calculated as the suprathreshold area divided by the total measured area. Relative fluorescence intensity was calculated as the mean suprathreshold fluorescence intensity after background subtraction. To minimize variability in immunostaining and image acquisition between staining batches, the fluorescence intensity of each *Bmp4* cKO section was normalized to that of the corresponding *Bmp4^fl/fl^* section processed in the same staining batch and imaged under identical acquisition settings.

For three-dimensional reconstruction of the basal forebrain, free-floating 50-μm sections were immunostained with an anti-ChAT antibody and visualized using nickel-enhanced DAB with Nissl counterstaining. ChAT-positive neurons were plotted using Neurolucida (MBF Bioscience, USA), and section contours were aligned to reconstruct the basal forebrain and neuronal distribution in three dimensions. Soma coordinates were rendered using Neurolucida Explorer (MBF Bioscience, USA).

For axonal morphology and varicosity analysis, AAV-injected mice expressing EGFP in cholinergic neurons were perfusion-fixed, and 50-μm free-floating coronal sections were prepared and co-immunostained with anti-GFP and anti-ChAT antibodies.

Neocortical sections were subsequently counterstained with thionin to visualize Nissl substance. Cholinergic axons were reconstructed using Neurolucida (MBF Bioscience, USA), and the boundaries of neocortical layers I–VI were determined from the Nissl-stained cytoarchitecture. Axon length and the numbers of branch points and varicosities were quantified for each neocortical layer. Axonal varicosities were quantified as morphological proxies for putative *en passant* boutons. Varicosity density was calculated as the number of varicosities divided by the reconstructed axon length within each neocortical layer.

### Statistical Analysis

Statistical analyses were performed using unpaired two-tailed Student’s *t*-tests, one-way ANOVA followed by the Holm–Šídák multiple-comparisons test, two-way ANOVA followed by the Holm–Šídák multiple-comparisons test, repeated-measures two-way ANOVA, or the Kolmogorov–Smirnov test, as appropriate for each dataset.

Non-normalized measurements were analyzed by two-way ANOVA with genotype and neocortical layer as factors, and both main effects and the genotype × neocortical layer interaction were evaluated (Figs. 4B, 6E, and 6F). For these analyses, Holm–Šídák-adjusted post hoc comparisons were performed between genotypes within each neocortical layer. Measurements normalized to an internal reference were analyzed by one-way ANOVA followed by the Holm–Šídák multiple-comparisons test because the normalization procedure altered the data structure such that the original factorial design was no longer applicable. This included normalization to the mean fluorescence intensity of VAChT-negative neurons within each image (Fig. 2F) and normalization to the fluorescence intensity of the corresponding *Bmp4^fl/fl^*section processed in the same staining batch and imaged under identical acquisition settings (Fig. 4C). Body weight was analyzed by repeated-measures two-way ANOVA with genotype and age as factors (Fig. 1E). For repeated-measures ANOVA, the Greenhouse–Geisser correction was applied when the assumption of sphericity was violated. The rostrocaudal distribution of cholinergic somata was compared using the Kolmogorov–Smirnov test (Fig. 5D). Data were assumed to follow a normal distribution for parametric analyses and are presented as mean ± standard error of the mean (SEM). All statistical analyses were performed using OriginPro (LightStone, Japan). Statistical significance was defined as *p* < 0.05. Statistical results are reported in the Results, with superscript lowercase letters referring to the corresponding entries in Table 3. Additional statistical information, including effect sizes, interval estimates, and sample sizes, is provided in Table 3 and the corresponding figure legends.

**Figure 1.**
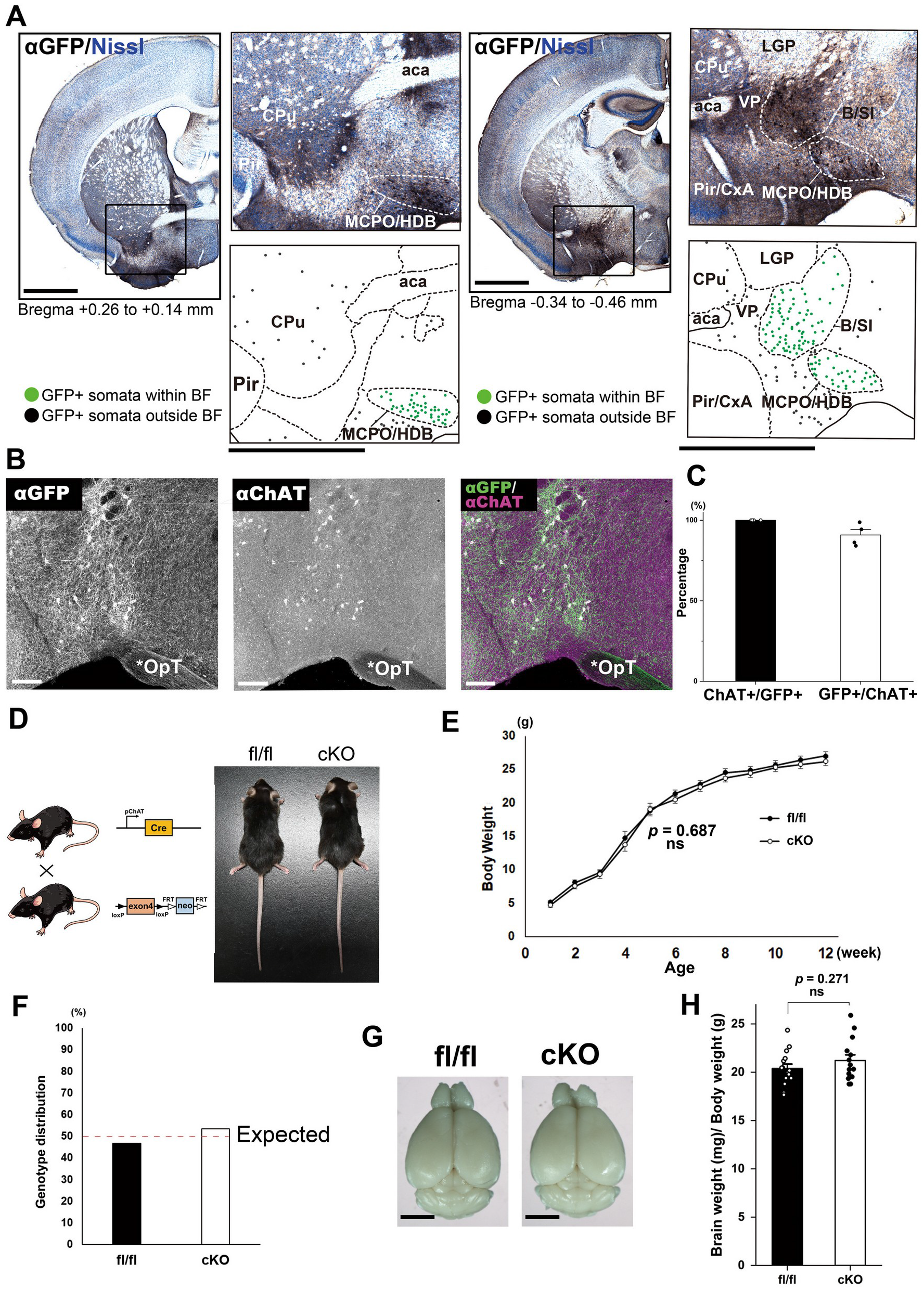
Generation of *Bmp4* cKO mice. (A) Representative coronal brain sections from mT/mG; ChAT-Cre mice, in which GFP-positive cells were visualized by DAB immunohistochemistry with Nissl counterstaining. Rostral (left; approximately +0.26 to +0.14 mm from bregma) and caudal (right; approximately −0.34 to −0.46 mm from bregma) levels of the basal forebrain are shown, based on Paxinos and Franklin (2001). At each rostrocaudal level, the boxed region in the low-magnification image is shown at a higher magnification (upper right), and the corresponding anatomical outline and positions of GFP-positive neuronal somata are plotted below (lower right). Green dots indicate GFP-positive neuronal somata within the outlined basal forebrain regions, whereas black dots indicate GFP-positive neuronal somata outside these regions. Abbreviations: aca, anterior commissure; BF, basal forebrain; B/SI, nucleus basalis of Meynert/substantia innominata; CPu, caudate–putamen; CxA, cortex–amygdala transition zone; HDB, horizontal limb of the diagonal band of Broca; LGP, lateral globus pallidus; MCPO, magnocellular preoptic nucleus; Pir, piriform cortex; VP, ventral pallidum. Scale bars, 1 mm in the low- and high-magnification images and corresponding anatomical plots. (B) Representative image of the basal forebrain from mT/mG; ChAT-Cre mice immunostained for GFP and ChAT. The asterisk indicates the optic tract (OpT). Scale bar, 50 μm. (C) Quantification of colocalization between GFP and ChAT in the basal forebrain of mT/mG; ChAT-Cre mice. Data are presented as mean ± SEM. Colocalization was evaluated as the percentage of ChAT-positive neurons that were GFP-positive (ChAT+/GFP+; left) and the percentage of GFP-positive cells that were ChAT-positive (GFP+/ChAT+; right). *n* = 4 coronal sections from one mT/mG; ChAT-Cre mouse; bilateral basal forebrain regions were analyzed in each section. (D) Breeding strategy for generating *Bmp4* cKO mice by crossing ChAT-Cre and *Bmp4* floxed mouse lines (left). Representative images of adult *Bmp4^fl/fl^* and *Bmp4* cKO mice are shown on the right. *Bmp4^fl/fl^*littermates lacking the ChAT-Cre transgene served as controls, whereas *Bmp4^fl/fl^*; ChAT-Cre littermates served as *Bmp4* cKO mice. (E) Body weight progression of *Bmp4^fl/fl^* and *Bmp4* cKO mice during the first 3 months of life. *n* = 9 *Bmp4^fl/fl^* mice and *n* = 11 *Bmp4* cKO mice. Data are presented as mean ± SEM. Statistical analysis was performed using repeated-measures two-way ANOVA; ns, not significant. See Table 3 for detailed statistical information. (F) Genotype distribution of offspring obtained from *Bmp4^fl/fl^*; ChAT-Cre × *Bmp4^fl/fl^* crosses. Because the ChAT-Cre transgene is hemizygous, *Bmp4^fl/fl^* and *Bmp4^fl/fl^*; ChAT-Cre offspring are expected at a 1:1 ratio (red dotted line). The observed numbers were 48 *Bmp4^fl/fl^* and 55 *Bmp4* cKO offspring. (G) Representative images of dissected whole brains from adult *Bmp4^fl/fl^* and *Bmp4* cKO mice. Scale bar, 3 mm. (H) Brain weight normalized to body weight in *Bmp4^fl/fl^* and *Bmp4* cKO mice. *n* = 15 *Bmp4^fl/fl^*mice and *n* = 14 *Bmp4* cKO mice. Data are presented as mean ± SEM. Statistical analysis was performed using an unpaired two-tailed Student’s *t*-test; ns, not significant. See Table 3 for detailed statistical information.

**Table 3.** Summary of statistical analyses.

| Dataset /<br>Comparison | Sample and analysis | Effect size | 95% CI (unadjusted) | Results |
|---|---|---|---|---|
| a<br>Fig. 1E<br>Body weight progression | *n*: fl/fl = 9 mice; cKO = 11 mice<br>Data structure: Normal distribution<br>Statistical test: Repeated-measures two-way ANOVA<br>Post hoc: — | Genotype: *F*(1, 18) = 0.168, partial η² = 0.009<br>Age: *F*(2.71, 48.74) = 959.37, partial η² = 0.982<br>Genotype × age: *F*(2.71, 48.74) = 0.623, partial η² = 0.033<br>Mean difference (cKO − fl/fl) = −0.31 g | Mean difference: [−1.90, 1.28] g | no significant main effect of genotype (*p* = 0.687)<br>significant main effect of age (*p* < 0.0001)<br>no significant genotype × age interaction (*p* = 0.588) |
| b<br>Fig. 1H<br>Brain weight/body weight | *n*: fl/fl = 15 mice; cKO = 14 mice<br>Data structure: Normal distribution<br>Statistical test: Unpaired two-tailed Student’s *t*-test<br>Post hoc: — | *t*(27) = −1.124<br>Cohen’s *d* (cKO − fl/fl) = 0.418<br>Mean difference (cKO − fl/fl) = 0.831 mg/g | Mean difference (cKO − fl/fl): [−0.686, 2.349] mg/g | no significant difference (*p* = 0.271) |
| c<br>Fig. 2C<br>VGluT1 area fraction | *n*: 5 regions from 3 mice/group<br>Data structure: Normal distribution<br>Statistical test: Unpaired two-tailed Student’s *t*-test<br>Post hoc: — | *t*(8) = 0.910<br>Cohen’s *d* (cKO − fl/fl) = −0.576<br>Mean difference (cKO − fl/fl) = −0.060 | Mean difference (cKO − fl/fl): [−0.211, 0.091] | no significant difference (*p* = 0.389) |
| d<br>Fig. 2C<br>VGAT area fraction | *n*: 7 regions from 4 mice/group<br>Data structure: Normal distribution<br>Statistical test: Unpaired two-tailed Student’s *t*-test<br>Post hoc: — | *t*(12) = 0.113<br>Cohen’s *d* (cKO − fl/fl) = −0.061<br>Mean difference (cKO − fl/fl) = −0.0036 | Mean difference (cKO − fl/fl): [−0.0724, 0.0652] | no significant difference (*p* = 0.912) |
| e<br>Fig. 2F<br>pSmad1/5 fluorescence intensity | *n*: 7 regions from 3 mice/group<br>Data structure: Normalized data<br>Statistical test: One-way ANOVA<br>Post hoc: Holm–Šídák | *F*(3, 24) = 40.285, η² = 0.834<br>fl/fl, VAChT(−) − VAChT(+): mean difference = −2.553<br>cKO, VAChT(−) − VAChT(+): mean difference = 0.170 | fl/fl, VAChT(−) − VAChT(+): [−3.154, −1.952]<br>cKO, VAChT(−) − VAChT(+): [−0.431, 0.771] | significant difference among groups (*p* < 0.0001)<br>fl/fl, VAChT(−) vs VAChT(+): *p* < 0.0001<br>cKO, VAChT(−) vs VAChT(+): *p* = 0.566 |
| f<br>Fig. 4B<br>VAChT-positive area fraction across neocortical layers I–VI | *n*: 9 regions from 3 mice/group<br>Data structure: Normal distribution<br>Statistical test: Two-way ANOVA<br>Post hoc: Holm–Šídák | Genotype: *F*(1, 80) = 87.156, partial η² = 0.521<br>Neocortical layer: *F*(4, 80) = 4.098, partial η² = 0.170<br>Genotype × neocortical layer: *F*(4, 80) = 6.464, partial η² = 0.244<br>Mean differences (cKO − fl/fl): I, 0.06989; II/III, 0.04895; IV, 0.00836; V, 0.04194; VI, 0.02423 | Mean differences (cKO − fl/fl): I, [0.0323, 0.1075]; II/III, [0.0358, 0.0621]; IV, [−0.00085, 0.01757]; V, [0.0318, 0.0521]; VI, [0.0118, 0.0367] | significant main effect of genotype (*p* < 0.0001)<br>significant main effect of neocortical layer (*p* = 0.00451)<br>significant genotype × neocortical layer interaction (*p* = 0.000147)<br>Holm–Šídák post hoc comparisons (cKO vs fl/fl within each layer): I, *p* = 0.002; II/III, *p* < 0.0001; IV, *p* = 0.072; V, *p* < 0.0001; VI, *p* = 0.002. |
| g<br>Fig. 4C<br>Relative VAChT fluorescence intensity across neocortical layers I–VI | *n*: 9 regions from 3 mice/group<br>Data structure: Normalized data<br>Statistical test: One-way ANOVA<br>Post hoc: Holm–Šídák | *F*(9, 80) = 5.765, η² = 0.393<br>Mean differences (cKO − fl/fl): I, 1.154; II/III, 1.093; IV, 1.726; V, 1.309; VI, 1.594 | Mean differences (cKO − fl/fl): I, [0.278, 2.031]; II/III, [0.216, 1.969]; IV, [0.850, 2.603]; V, [0.432, 2.185]; VI, [0.718, 2.471] | significant difference among groups (*p* < 0.0001)<br>Holm–Šídák post hoc comparisons (cKO vs fl/fl within each layer): I, *p* = 0.0104; II/III, *p* = 0.015; IV, *p* = 0.0002; V, *p* = 0.004; VI, *p* = 0.0005. |
| h<br>Fig. 5C<br>Number of ChAT-positive neuronal somata | *n*: fl/fl = 3 mice; cKO = 3 mice<br>Data structure: Normal distribution<br>Statistical test: Unpaired two-tailed Student’s *t*-test<br>Post hoc: — | *t*(4) = 0.907<br>Cohen’s *d* (cKO − fl/fl) = −0.740<br>Mean difference (cKO − fl/fl) = −1487.33 neurons | Mean difference (cKO − fl/fl): [−6041.1, 3066.4] neurons | no significant difference (*p* = 0.416) |
| i<br>Fig. 5D<br>Rostrocaudal distribution of ChAT-positive somata | *n*: fl/fl = 3 mice; cKO = 3 mice<br>Data structure: Distribution<br>Statistical test: Two-sample Kolmogorov–Smirnov test<br>Post hoc: — | *D* = 0.12 | Not applicable | no significant difference (*p* = 0.996) |
| j<br>Fig. 6E<br>Axonal branch points per neocortical layer | *n*: *Bmp4*<sup>+/+</sup>; ChAT-Cre = 4 mice; *Bmp4* cKO = 3 mice<br>Data structure: Normal distribution<br>Statistical test: Two-way ANOVA<br>Post hoc: Holm–Šídák | Genotype: *F*(1, 96) = 7.484, partial η² = 0.072<br>Neocortical layer: *F*(4, 96) = 2.914, partial η² = 0.108<br>Genotype × neocortical layer: *F*(4, 96) = 3.071, partial η² = 0.113<br>Mean differences (cKO − control): I, 0.400; II/III, 2.356; IV, 0.111; V, 4.392; VI, −0.455 | Mean differences (cKO − control): I, [−0.891, 1.691]; II/III, [−1.361, 6.073]; IV, [−1.140, 1.362]; V, [2.315, 6.468]; VI, [−1.724, 0.815] | significant main effect of genotype (*p* = 0.00742)<br>significant main effect of neocortical layer (*p* = 0.02532)<br>significant genotype × neocortical layer interaction (*p* = 0.01993)<br>Holm–Šídák post hoc comparisons (cKO vs control within each layer): I, *p* = 0.846; II/III, *p* = 0.594; IV, *p* = 0.854; V, *p* = 0.0026; VI, *p* = 0.846. |
| k<br>Fig. 6F<br>Varicosity density along cholinergic axons | *n*: *Bmp4*<sup>+/+</sup>; ChAT-Cre = 4 mice; *Bmp4* cKO = 3 mice<br>Data structure: Normal distribution<br>Statistical test: Two-way ANOVA<br>Post hoc: Holm–Šídák | Genotype: *F*(1, 88) = 35.456, partial η² = 0.287<br>Neocortical layer: *F*(4, 88) = 0.366, partial η² = 0.016<br>Genotype × neocortical layer: *F*(4, 88) = 0.770, partial η² = 0.034<br>Mean differences (cKO − control): I, 0.116; II/III, 0.221; IV, 0.109; V, 0.134; VI, 0.197 | Mean differences (cKO − control): I, [−0.023, 0.254]; II/III, [0.096, 0.346]; IV, [−0.029, 0.247]; V, [0.025, 0.243]; VI, [0.101, 0.292] | significant main effect of genotype (*p* < 0.0001)<br>no significant main effect of neocortical layer (*p* = 0.832)<br>no significant genotype × neocortical layer interaction (*p* = 0.548)<br>Holm–Šídák post hoc comparisons (cKO vs control within each layer): I, *p* = 0.179; II/III, *p* = 0.006; IV, *p* = 0.179; V, *p* = 0.055; VI, *p* = 0.0021. |

Notes: Unless otherwise specified, mean differences and Cohen’s *d* for genotype comparisons are expressed as cKO − control (*Bmp4*<sup>*fl/fl*</sup> or *Bmp4*<sup>+/+</sup>; ChAT-Cre). For unpaired *t*-tests, the reported *t* statistics were calculated as control − cKO; therefore, their signs are opposite to those of the corresponding mean differences and Cohen’s *d*. For Fig. 2F, mean differences are expressed as VAChT(−) − VAChT(+). Confidence intervals for post hoc comparisons are unadjusted, whereas post hoc *p* values are Holm–Šídák-adjusted.

## Results

### Generation of Cholinergic Neuron-specific Bmp4 Knockout Mice

To selectively delete *Bmp4* in cholinergic neurons, we used ChAT-Cre transgenic mice, which express Cre recombinase in most cholinergic neurons throughout the central nervous system (Gong et al., 2007). To verify the distribution of Cre activity, ChAT-Cre mice were crossed with mT/mG reporter mice (Muzumdar et al., 2007). In this reporter line, membrane-targeted tdTomato (mT) is constitutively expressed and is replaced by membrane-targeted EGFP (mG) following Cre-mediated recombination. GFP immunoreactivity was visualized using DAB staining. Consistent with the reported distribution of ChAT-Cre expression (Gong et al., 2007), GFP-positive neurons were observed in cholinergic neuron-rich regions, including the striatum, medial septum, diagonal band of Broca, ventral pallidum, and nucleus basalis of Meynert (Fig. 1A).

Double immunofluorescence staining with an anti-ChAT antibody showed that all ChAT-positive neurons in the basal forebrain were GFP-positive and that 84–99% of GFP-positive neurons were ChAT-positive (Fig. 1B, C). These results indicate that Cre recombination occurred in all cholinergic neurons examined in the basal forebrain and was largely restricted to them.

*Bmp4* cKO mice were generated by crossing ChAT-Cre transgenic mice with *Bmp4* floxed mice (Liu et al., 2004) as outlined in Fig. 1D. *Bmp4^fl/fl^*; ChAT-Cre littermates were used as *Bmp4* cKO mice, whereas *Bmp4^fl/fl^* littermates lacking the ChAT-Cre transgene served as controls. Body weight was similar between *Bmp4^fl/fl^* and *Bmp4* cKO mice during the first 3 months of life (genotype: *F*(1, 18) = 0.168, *p* = 0.687; genotype × age: *F*(2.71, 48.74) = 0.623, *p* = 0.588ᵃ; Fig. 1E). The genotype distribution of offspring was consistent with the expected 1:1 ratio (48 *Bmp4^fl/fl^* and 55 *Bmp4* cKO; Fig. 1F). Likewise, brain weight relative to body weight at 3 months of age did not differ between *Bmp4^fl/fl^*and *Bmp4* cKO mice (*t*(27) = −1.124, *p* = 0.271ᵇ; Fig. 1G, H).

Cholinergic neuron-specific deletion of *Bmp4* did not cause overt abnormalities in viability, postnatal growth, or gross brain morphology.

### Bmp4 Deletion Does Not Alter Excitatory or Inhibitory Presynaptic Terminals in Neocortical Layer II/III

Proper neural circuit function depends on a balance between excitatory and inhibitory synaptic inputs (E/I balance). A recent study demonstrated that BMP2 secreted from parvalbumin-positive interneurons contributes to the regulation of E/I balance (Okur et al., 2024). Because BMP2 and BMP4 belong to the same BMP2/4 subfamily and activate common downstream signaling pathways, we first asked whether BMP4 derived from cholinergic neurons affects the abundance of excitatory and inhibitory presynaptic terminals in the neocortex. We quantified VGluT1-positive glutamatergic and VGAT-positive GABAergic presynaptic terminals in layer II/III of the primary somatosensory cortex, where cholinergic innervation is rich (Ballinger et al., 2016; Fig. 2A, B). Layer IV was not analyzed because the barrel field receives dense thalamocortical inputs with a distinct synaptic organization (Woolsey and Van der Loos, 1970; Petersen, 2007; Erzurumlu and Gaspar, 2012).

**Figure 2.**
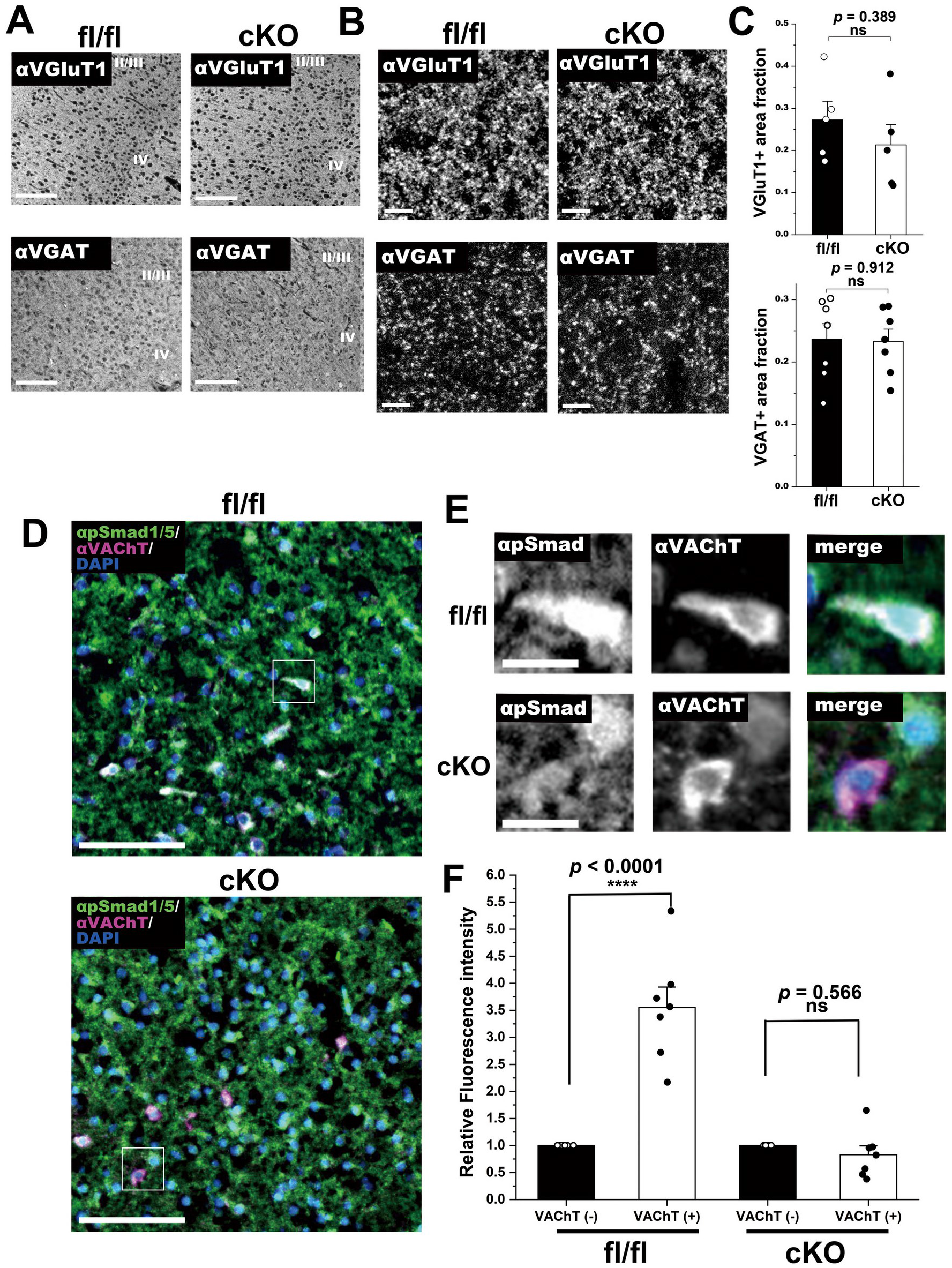
Cortical excitatory and inhibitory presynaptic terminals and BMP–Smad signaling in basal forebrain cholinergic neurons of *Bmp4* cKO mice. (A) Representative low-magnification fluorescence images showing VGluT1 and VGAT immunostaining in the primary somatosensory cortex from *Bmp4^fl/fl^* and *Bmp4* cKO mice. Scale bar, 100 μm. (B) Higher-magnification images showing VGluT1 and VGAT immunostaining in layer II/III of the primary somatosensory cortex. Scale bar, 5 μm. (C) Quantification of the area fraction of VGluT1-positive and VGAT-positive signals in layer II/III of the primary somatosensory cortex. Data are presented as mean ± SEM. For VGluT1, *n* = 5 regions from 3 *Bmp4^fl/fl^* mice and *n* = 5 regions from 3 *Bmp4* cKO mice. For VGAT, *n* = 7 regions from 4 *Bmp4^fl/fl^* mice and *n* = 7 regions from 4 *Bmp4* cKO mice. Statistical analysis was performed using an unpaired two-tailed Student’s *t*-test; ns, not significant. See Table 3 for detailed statistical information. (D) Representative fluorescence images showing pSmad1/5 (green) and VAChT (magenta) immunostaining with DAPI nuclear staining (blue) in the basal forebrain of *Bmp4^fl/fl^* and *Bmp4* cKO mice at postnatal days 4–5. Scale bar, 20 μm. (E) Higher-magnification images of the boxed regions in (D), showing pSmad1/5 (green) and VAChT (magenta), with DAPI nuclear staining (blue) in the merged images. Scale bar, 5 μm. (F) Quantification of pSmad1/5 fluorescence intensity in VAChT-positive and VAChT-negative neurons in the basal forebrain. pSmad1/5 fluorescence intensity was normalized to the mean fluorescence intensity of VAChT-negative neurons within each image. Data are presented as mean ± SEM. In *Bmp4^fl/fl^* mice, 94 VAChT-negative and 133 VAChT-positive neurons were analyzed from 7 regions obtained from 3 mice. In *Bmp4* cKO mice, 76 VAChT-negative and 83 VAChT-positive neurons were analyzed from 7 regions obtained from 3 mice. Statistical analysis was performed using one-way ANOVA. \*\*\*\**p* < 0.0001 by Holm–Šídák-adjusted post hoc comparisons; ns, not significant. See Table 3 for detailed statistical information.

The area fraction of VGluT1-positive terminals did not differ between *Bmp4^fl/fl^* and *Bmp4* cKO mice (*t*(8) = 0.910, *p* = 0.389ᶜ; Fig. 2C), nor did that of VGAT-positive terminals (*t*(12) = 0.113, *p* = 0.912ᵈ; Fig. 2C). *Bmp4* deletion in cholinergic neurons did not detectably change the area fraction of excitatory or inhibitory presynaptic terminals in neocortical layer II/III. We next focused on the cholinergic neurons themselves.

### Bmp4 cKO Mice Lack Elevated Smad1/5 Phosphorylation in Basal Forebrain Cholinergic Neurons

In the rat, cholinergic projections from the basal forebrain undergo a marked increase in terminal density between postnatal days 4–5 and 7–8 (Mechawar and Descarries, 2001). We therefore hypothesized that BMP4 signaling is active in cholinergic neurons during this developmental period. To test this hypothesis, we measured Smad1/5 phosphorylation as a readout of canonical BMP signaling (Miyazono et al., 2010; Katagiri and Watabe, 2016) in brain sections obtained from mice at postnatal days 4–5. In *Bmp4^fl/fl^* mice, pSmad1/5 immunoreactivity was detected in a subset of basal forebrain neurons and was significantly higher in VAChT-positive cholinergic neurons than in VAChT-negative neurons (*p* < 0.0001ᵉ; Fig. 2D–F). In *Bmp4* cKO mice, pSmad1/5 fluorescence intensity did not differ between VAChT-positive and VAChT-negative neurons (*p* = 0.566ᵉ; Fig. 2F). Thus, BMP–Smad signaling is enriched in basal forebrain cholinergic neurons during the period of rapid cholinergic terminal growth, and this enrichment depends on BMP4 expressed in cholinergic neurons.

### Bmp4 Deletion Increases Cholinergic Axon Terminals in Projection Targets

BMP4 secreted from axon terminals of hippocampal glutamatergic neurons locally suppresses presynaptic terminal formation (Higashi et al., 2018), and BMP–Smad signaling was enriched in cholinergic neurons during the period of rapid cholinergic terminal growth (Fig. 2D–F). These observations led us to hypothesize that deletion of *Bmp4* from cholinergic neurons increases their axon terminals within projection targets. To test this hypothesis, we visualized cholinergic axon terminals by immunostaining for VAChT in the neocortex, hippocampus, and amygdala, where cholinergic terminals derive predominantly from basal forebrain projection neurons. Cholinergic innervation appeared denser in *Bmp4* cKO mice than in *Bmp4^fl/fl^* mice in all three regions (Fig. 3A–C).

**Figure 3.**
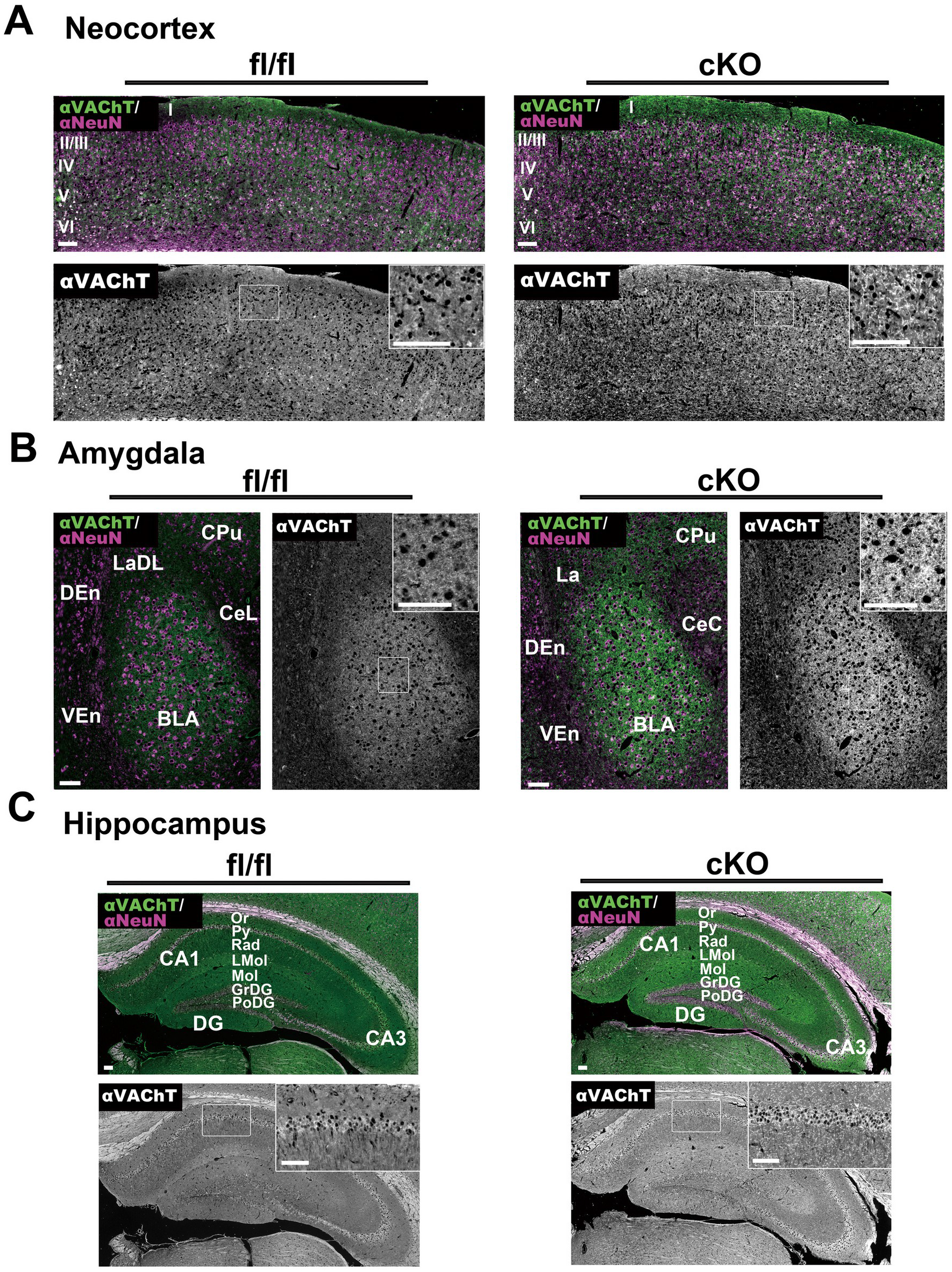
*Bmp4* deletion increases cholinergic axon terminals in projection targets. Representative immunofluorescence images showing VAChT (green) and NeuN (magenta) in the neocortex (A), amygdala (B), and hippocampus (C) of *Bmp4^fl/fl^* and *Bmp4* cKO mice. Merged images show VAChT and NeuN immunofluorescence, whereas the corresponding VAChT immunoreactivity alone is shown in grayscale. White boxes in the grayscale images indicate regions shown at higher magnification in the insets. (A) Neocortical layers are indicated by Roman numerals: I, molecular layer; II/III, external granular and external pyramidal layers; IV, internal granular layer; V, internal pyramidal layer; VI, multiform layer. (B) Abbreviations: BLA, basolateral amygdaloid nucleus; CeC, capsular part of the central amygdaloid nucleus; CeL, lateral part of the central amygdaloid nucleus; CPu, caudate–putamen; DEn, dorsal endopiriform nucleus; La, lateral amygdaloid nucleus; LaDL, dorsolateral part of the lateral amygdaloid nucleus; VEn, ventral endopiriform nucleus. (C) Abbreviations: CA1–CA3, Ammon’s horn regions; DG, dentate gyrus; GrDG, granule cell layer of the dentate gyrus; LMol, stratum lacunosum-moleculare; Mol, molecular layer; Or, stratum oriens; PoDG, polymorphic layer of the dentate gyrus; Py, pyramidal cell layer; Rad, stratum radiatum. Anatomical nomenclature is based on Paxinos and Franklin (2001). Scale bars, 100 μm.

To quantify this increase, we measured VAChT-positive terminals across layers I–VI of the primary somatosensory cortex. Neocortical layers were identified by NeuN immunostaining and DAPI labeling (Fig. 4A). Layer IV was identified by the characteristic barrel structures receiving thalamocortical inputs, whereas layer V was distinguished by the presence of large pyramidal neurons. Two-way ANOVA of the VAChT-positive terminal area fraction revealed significant main effects of genotype (*F*(1, 80) = 87.156, *p* < 0.0001) and neocortical layer (*F*(4, 80) = 4.098, *p* = 0.00451), as well as a significant genotype × neocortical layer interaction (*F*(4, 80) = 6.464, *p* = 0.000147ᶠ) (Fig. 4B). The area fraction in *Bmp4* cKO mice was 1.91-to 3.58-fold that in *Bmp4^fl/fl^* mice in layers I, II/III, V, and VI, and Holm–Šídák post hoc comparisons showed significant increases in layers I (*p* = 0.002), II/III (*p* < 0.0001), V (*p* < 0.0001), and VI (*p* = 0.002), but not in layer IV (*p* = 0.072ᶠ). In both genotypes, VAChT-positive axons remained sparse within the barrel structures of layer IV (Fig. 4A). Relative VAChT fluorescence intensity in *Bmp4* cKO mice was also 2.09-to 2.73-fold the *Bmp4^fl/fl^* level in all neocortical layers, including layer IV (one-way ANOVA: *F*(9, 80) = 5.765, *p* < 0.0001; Holm–Šídák post hoc: layer I, *p* = 0.0104; layer II/III, *p* = 0.015; layer IV, *p* = 0.0002; layer V, *p* = 0.004; layer VI, *p* = 0.0005ᵍ; Fig. 4C). In layer IV, VAChT immunoreactivity increased without a detectable expansion of the VAChT-positive area.

**Figure 4.**
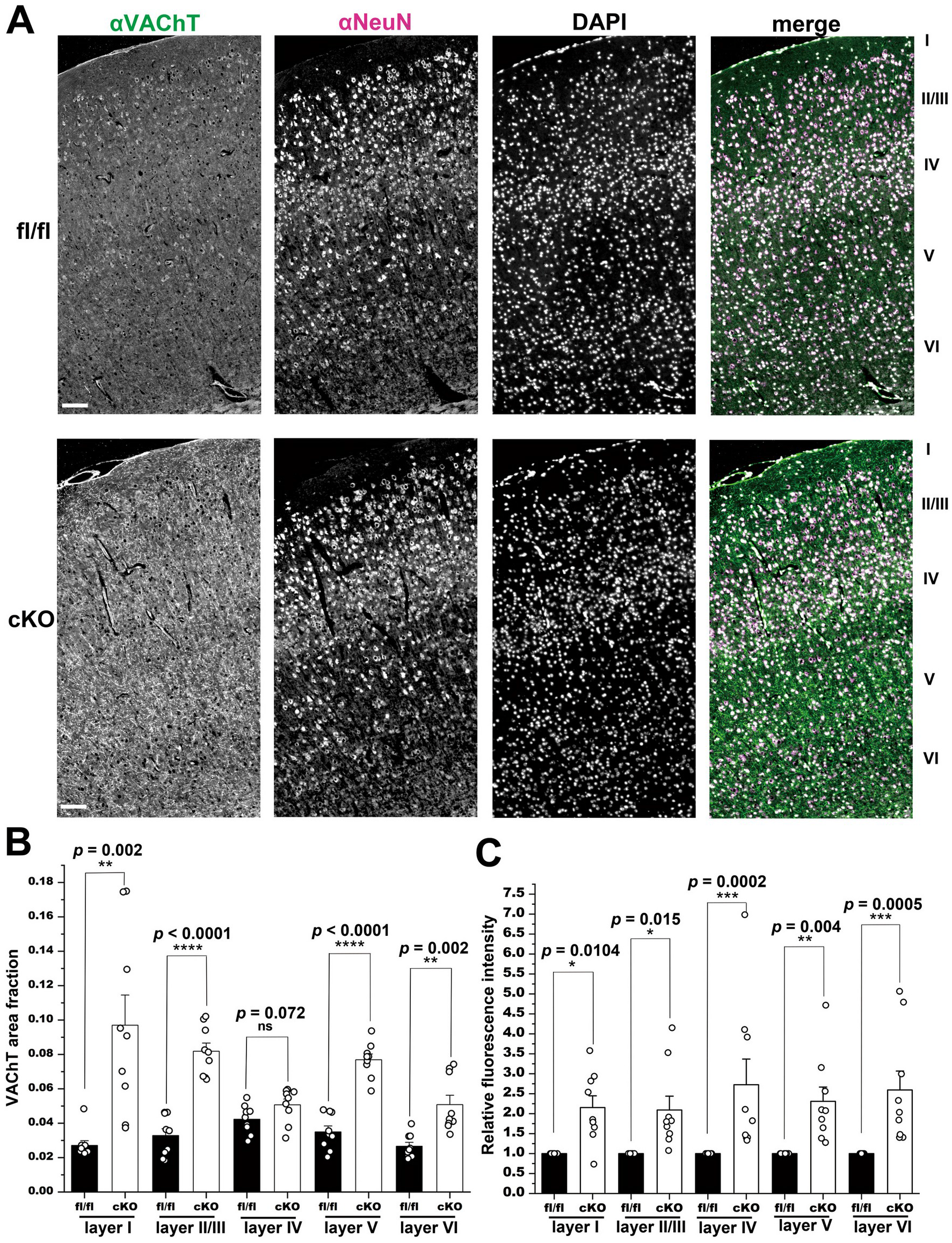
*Bmp4* deletion increases cholinergic innervation across neocortical layers. (A) Representative immunofluorescence images showing VAChT (green) and NeuN (magenta), together with DAPI nuclear staining (gray), in coronal sections of the primary somatosensory cortex from *Bmp4^fl/fl^* and *Bmp4* cKO mice. Neocortical layers I–VI are indicated. Scale bar, 100 μm. (B) Quantification of VAChT-positive terminal area fraction across neocortical layers I– VI of the primary somatosensory cortex. Data are presented as mean ± SEM. VAChT-positive terminal area fraction was calculated as the suprathreshold area divided by the total measured area. Data were analyzed by two-way ANOVA with genotype and neocortical layer as factors. Significant main effects of genotype and neocortical layer, as well as a significant genotype × neocortical layer interaction, were detected. \*\**p* < 0.01, \*\*\*\**p* < 0.0001 by Holm–Šídák-adjusted post hoc comparisons between genotypes within each neocortical layer; ns, not significant. *Bmp4^fl/fl^*, *n* = 9 regions from 3 mice; *Bmp4* cKO, *n* = 9 regions from 3 mice. See Table 3 for detailed statistical information. (C) Quantification of relative VAChT fluorescence intensity across neocortical layers I– VI of the primary somatosensory cortex. Data are presented as mean ± SEM. Fluorescence intensity in each *Bmp4* cKO section was normalized to that of the corresponding *Bmp4^fl/fl^* section processed in the same staining batch and imaged under identical acquisition settings. Data were analyzed by one-way ANOVA. \**p* < 0.05, \*\**p* < 0.01, \*\*\**p <* 0.001 by Holm–Šídák-adjusted post hoc comparisons between genotypes within each neocortical layer. *Bmp4^fl/fl^*, *n* = 9 regions from 3 mice; *Bmp4* cKO, *n* = 9 regions from 3 mice. See Table 3 for detailed statistical information.

### Bmp4 Deletion Does Not Alter the Number or Rostrocaudal Distribution of Cholinergic Neurons in the Basal Forebrain

How does the increase in cholinergic terminals arise? We considered two possibilities: (1) an increase in the number of basal forebrain cholinergic neurons or (2) an increase in axonal branching. As BMP4 regulates cell proliferation and differentiation in a variety of tissues (Miyazono et al., 2010; Katagiri and Watabe, 2016; Sadler, 2019; Alvarez et al., 2021), we first examined whether deletion of *Bmp4* affects the cholinergic neuron population in the basal forebrain.

Serial coronal sections were immunostained for ChAT and visualized using nickel-enhanced DAB with Nissl counterstaining. Based on the mouse brain atlas (Paxinos and Franklin, 2001), the medial septum (MS), vertical and horizontal limbs of the diagonal band of Broca (VDB and HDB), nucleus basalis of Meynert/substantia innominata (B/SI), and ventral pallidum (VP) were identified, and ChAT-positive somata were counted in each region (Fig. 5A).

**Figure 5.**
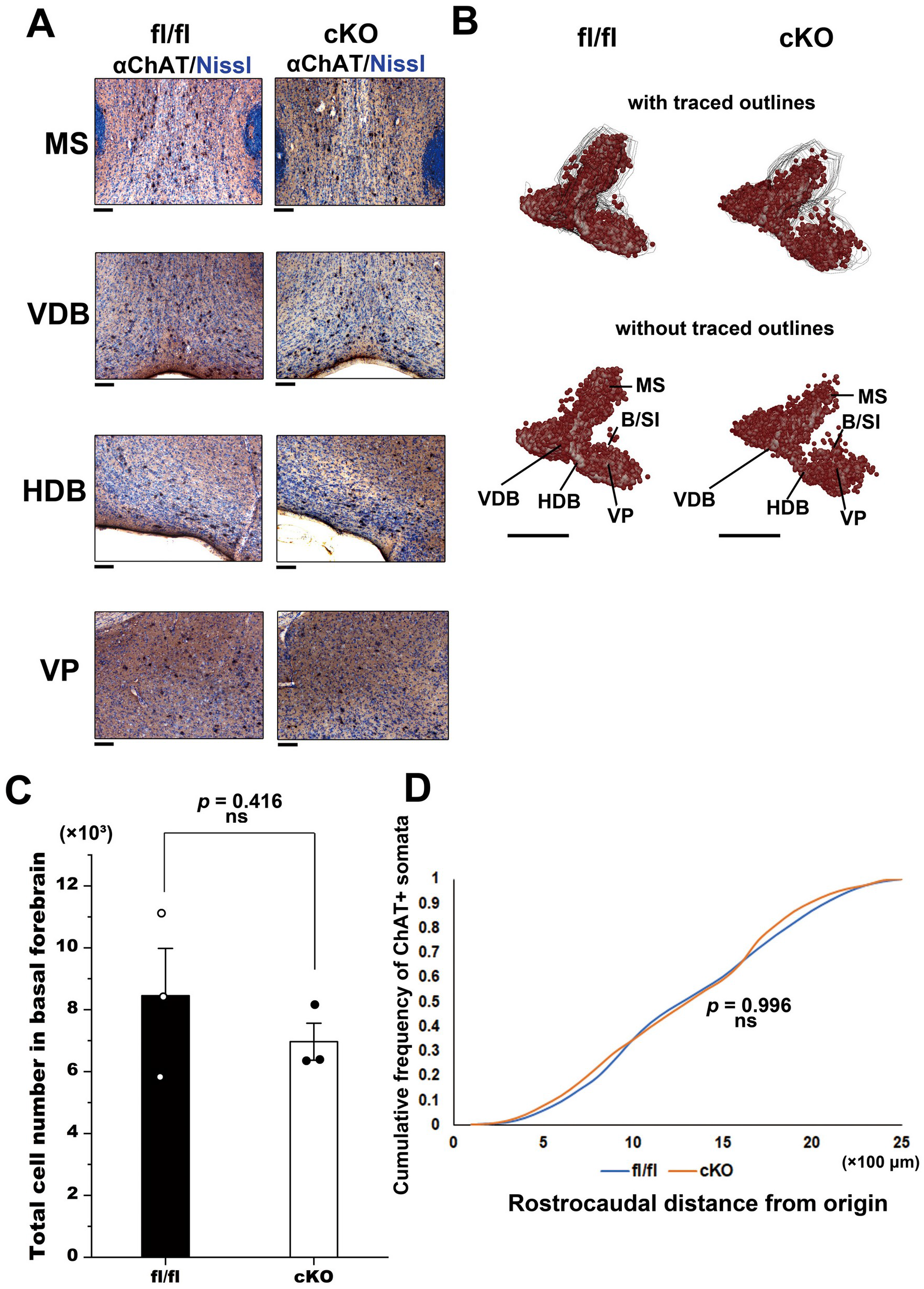
*Bmp4* deletion does not alter the number or rostrocaudal distribution of basal forebrain cholinergic neurons. (A) Representative images showing ChAT-positive neurons visualized by nickel-enhanced DAB immunohistochemistry with Nissl counterstaining in the medial septum (MS), vertical limb of the diagonal band of Broca (VDB), horizontal limb of the diagonal band of Broca (HDB), and ventral pallidum (VP). The nucleus basalis of Meynert/substantia innominata (B/SI) was also included in the regions used for cell counting and three-dimensional reconstruction (B–D). Scale bar, 100 μm. (B) Three-dimensional reconstruction of ChAT-positive neuronal somata in the basal forebrain from serial coronal sections. ChAT-positive somata were plotted and basal forebrain outlines were traced using Neurolucida, and the reconstructed neuronal distribution was rendered using Neurolucida Explorer. Upper panels show ChAT-positive somata together with the manually traced outlines of the basal forebrain in each coronal section, whereas lower panels show the same somata without the outlines. Scale bar, 1 mm. (C) Quantification of the number of ChAT-positive neuronal somata in the basal forebrain. No significant difference was detected between *Bmp4^fl/fl^* and *Bmp4* cKO mice. Data are presented as mean ± SEM. Statistical analysis was performed using an unpaired two-tailed Student’s *t*-test; ns, not significant. *Bmp4^fl/fl^*, *n* = 3 mice; *Bmp4* cKO, *n* = 3 mice. See Table 3 for detailed statistical information. (D) Cumulative frequency distributions of the rostrocaudal positions of ChAT-positive somata in the basal forebrain. The most rostral section containing ChAT-positive somata within the MS/VDB was defined as the origin for each mouse. No significant difference was detected between *Bmp4^fl/fl^* and *Bmp4* cKO mice. Statistical analysis was performed using the Kolmogorov–Smirnov test; ns, not significant. *Bmp4^fl/fl^*, *n* = 3 mice; *Bmp4* cKO, *n* = 3 mice. See Table 3 for detailed statistical information.

The three-dimensional distribution of ChAT-positive neurons was reconstructed using Neurolucida (Fig. 5B). The total number of basal forebrain cholinergic neurons did not differ between *Bmp4^fl/fl^* and *Bmp4* cKO mice (*t*(4) = 0.907, *p* = 0.416ʰ; Fig. 5C). Because basal forebrain cholinergic neurons are topographically organized along the rostrocaudal axis according to their projection targets (Zaborszky et al., 2012; Bloem et al., 2014), we next compared their rostrocaudal distribution. Using the most rostral section containing ChAT-positive neurons in the MS/VDB as the reference point, cumulative distributions of cholinergic neurons were generated for each genotype. No significant difference was detected between *Bmp4^fl/fl^* and *Bmp4* cKO mice (*D* = 0.12, *p* = 0.996ⁱ; Fig. 5D). The increase in cholinergic terminals is therefore unlikely to be explained by a change in the cholinergic neuron population.

### Bmp4 Deletion Increases Cholinergic Axon Ramification and Varicosity Density in the Neocortex

Finally, to test the second possibility, that *Bmp4* deletion increases axonal branching, we stereotaxically injected an AAV9-pCAG-FLEX-EGFP-WPRE vector into the rostral and caudal basal forebrain of *Bmp4^+/+^*; ChAT-Cre and *Bmp4* cKO mice to label cholinergic neurons (Fig. 6A). Immunostaining with anti-ChAT and anti-GFP antibodies confirmed that most GFP-expressing cells were ChAT-positive neurons (Fig. 6B).

**Figure 6.**
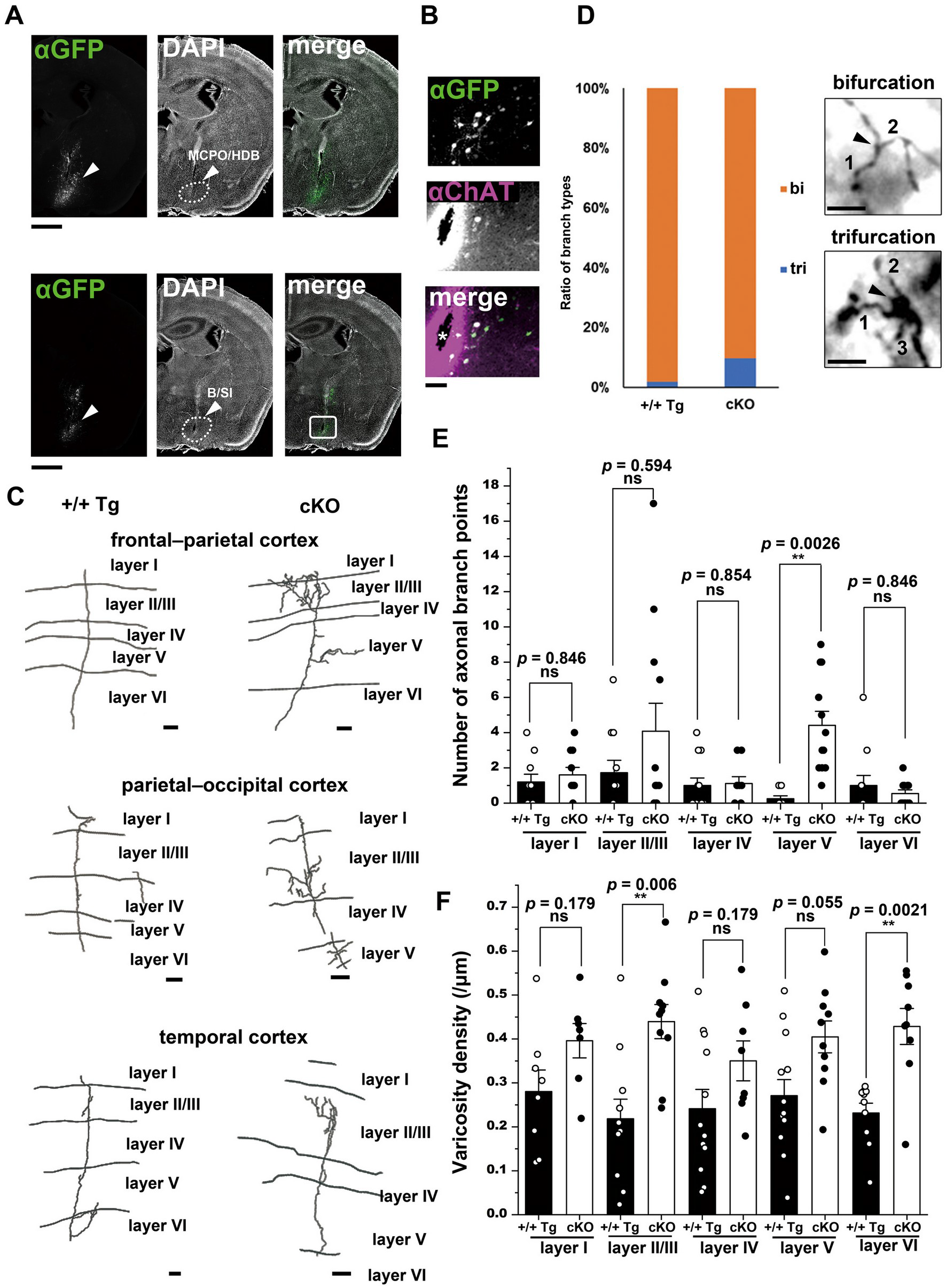
*Bmp4* deletion increases axonal ramification and varicosity density inneocortical cholinergic projections. (A) Representative coronal sections showing AAV injection sites in the rostral (upper) and caudal (lower) basal forebrain. GFP immunoreactivity is shown in green and DAPI nuclear staining in gray. Arrowheads indicate the injection sites, and dashed lines indicate the boundaries of the MCPO/HDB (upper) and B/SI (lower), delineated based on DAPI staining and Paxinos and Franklin (2001). B/SI, nucleus basalis of Meynert/substantia innominata; HDB, horizontal limb of the diagonal band of Broca; MCPO, magnocellular preoptic nucleus. Scale bar, 1 mm. (B) Higher-magnification image of the boxed region in the lower merged panel of (A), with GFP (green) and ChAT (magenta) immunoreactivity. The injection track is indicated by an asterisk. Scale bar, 100 μm. (C) Representative Neurolucida reconstructions of basal forebrain-derived cholinergic axons in the neocortex. Top, frontal–parietal cortex; middle, parietal–occipital cortex; bottom, temporal cortex. Scale bar, 20 μm. In C–F, +/+ Tg denotes *Bmp4^+/+^*; ChAT-Cre mice, whereas cKO denotes *Bmp4^fl/fl^*; ChAT-Cre mice. (D) Left, proportion of axonal branch points classified as bifurcations (bi) or trifurcations (tri). Right, representative examples of bifurcation and trifurcation morphologies. Arrowheads indicate branch points, and numbers indicate daughter branches. Scale bar, 5 μm. (E) Quantification of axonal branch points across neocortical layers I–VI. Data are presented as mean ± SEM. Data were analyzed by two-way ANOVA with genotype and neocortical layer as factors. Significant main effects of genotype and neocortical layer, as well as a significant genotype × neocortical layer interaction, were detected. \*\**p* < 0.01 by Holm–Šídák-adjusted post hoc comparisons between genotypes within each neocortical layer; ns, not significant. *Bmp4^+/+^*; ChAT-Cre mice: *n* = 4 mice (layer I, 10 axons; II/III, 11 axons; IV, 12 axons; V, 8 axons; VI, 11 axons). *Bmp4* cKO mice: *n* = 3 mice (layer I, 10 axons; II/III, 12 axons; IV, 9 axons; V, 12 axons; VI, 11 axons). See Table 3 for detailed statistical information. (F) Quantification of varicosity density along cholinergic axons across neocortical layers I–VI. Varicosity density is expressed as the number of varicosities per micrometer of axon length. Data are presented as mean ± SEM. Data were analyzed by two-way ANOVA with genotype and neocortical layer as factors. A significant main effect of genotype was detected, whereas the main effect of neocortical layer and the genotype × neocortical layer interaction were not significant. \*\**p* < 0.01 by Holm– Šídák-adjusted post hoc comparisons between genotypes within each neocortical layer; ns, not significant. *Bmp4^+/+^*; ChAT-Cre mice: *n* = 4 mice (layer I, 8 axons; II/III, 11 axons; IV, 12 axons; V, 13 axons; VI, 10 axons). *Bmp4* cKO mice: *n* = 3 mice (layer I, 7 axons; II/III, 10 axons; IV, 8 axons; V, 10 axons; VI, 9 axons). See Table 3 for detailed statistical information.

Labeled axons were reconstructed in the frontal–parietal, parietal–occipital, and temporal cortical areas (Fig. 6C). Most branch points were bifurcations, whereas a subset were trifurcations, with a higher proportion of trifurcations observed in *Bmp4* cKO mice (Fig. 6D).

Two-way ANOVA of axonal branch points revealed significant main effects of genotype (*F*(1, 96) = 7.484, *p* = 0.00742) and neocortical layer (*F*(4, 96) = 2.914, *p* = 0.02532), as well as a significant genotype × neocortical layer interaction (*F*(4, 96) = 3.071, *p* = 0.01993ʲ). Holm–Šídák post hoc comparisons showed a significant increase in branch points in layer V of *Bmp4* cKO mice (*p* = 0.0026ʲ; Fig. 6E). *Bmp4* deletion increased axonal ramification in a layer-dependent manner.

Two-way ANOVA of varicosity density revealed a significant main effect of genotype (*F*(1, 88) = 35.456, *p* < 0.0001), but neither a significant main effect of neocortical layer (*F*(4, 88) = 0.366, *p* = 0.832) nor a significant genotype × neocortical layer interaction (*F*(4, 88) = 0.770, *p* = 0.548ᵏ). Varicosity density was increased overall, without a layer-dependent difference (Fig. 6F). Together with the unchanged number and distribution of cholinergic neurons (Fig. 5), these results indicate that BMP4 restricts the ramification and varicosity density of basal forebrain cholinergic axons. e 3. Summary of statistical analyses.

## Discussion

### BMP4–Smad Signaling in Basal Forebrain Cholinergic Neurons

BMP4 is broadly expressed in the postnatal central nervous system, including the basal forebrain (Mikawa et al., 2006; Higashi et al., 2018). In this study, Smad1/5 phosphorylation was enriched in basal forebrain cholinergic neurons at postnatal days 4–5, and this enrichment depended on BMP4 expressed in these neurons (Fig. 2D–F).

This period coincides with the rapid increase in cholinergic terminals in the neocortex (Mechawar and Descarries, 2001). *Bmp4* cKO mice showed no overt abnormalities in viability, postnatal growth, or brain weight (Fig. 1). The axonal changes described below are thus unlikely to reflect general developmental defects.

### Basal Forebrain Cholinergic Neuron Number and Rostrocaudal Distribution in Bmp4 cKO Mice

Three-dimensional reconstruction and quantitative analysis of the basal forebrain showed no significant differences in cholinergic neuron number or rostrocaudal distribution between *Bmp4^fl/fl^* and *Bmp4* cKO mice (Fig. 5). We focused on the rostrocaudal axis because basal forebrain cholinergic neurons are topographically organized along this axis according to their cortical targets (Zaborszky et al., 2012; Bloem et al., 2014). Changes in neuron number or distribution are therefore unlikely to account for the increased cholinergic innervation in *Bmp4* cKO mice.

Previous studies have shown that BMP signaling can induce and maintain the cholinergic phenotype in the central nervous system (López-Coviella et al., 2000) and regulate the specification of dorsal interneurons during neural tube development (Alvarez et al., 2021). In the rat, basal forebrain cholinergic neurons are generated during embryonic development (Semba and Fibiger, 1988), whereas their cortical innervation expands rapidly during the first postnatal weeks (Mechawar and Descarries, 2001). In *Bmp4* cKO mice, the number and rostrocaudal distribution of basal forebrain cholinergic neurons were not detectably altered, whereas axonal ramification and varicosity density were increased in their neocortical projections. Together with the enrichment of Smad1/5 phosphorylation in cholinergic neurons at postnatal days 4–5, these findings support a role for BMP4 in the postnatal maturation of cholinergic projections. Because the onset of Cre-mediated recombination in the GM24Gsat ChAT-Cre line has not been determined, temporally controlled deletion of *Bmp4* will be needed to distinguish its embryonic, postnatal, and adult functions.

Cholinergic interneurons expressing vasoactive intestinal peptide have been reported in the neocortex (Granger et al., 2020). However, we did not detect ChAT- or VAChT-positive neuronal somata in the neocortex under our immunohistochemical conditions, possibly reflecting low expression levels of these markers in cortical interneurons.

Furthermore, AAV-mediated labeling showed that cholinergic axons originating from the basal forebrain had more branch points and a higher varicosity density in *Bmp4* cKO mice (Fig. 6). These findings support the interpretation that the increased cholinergic innervation in the neocortex predominantly reflects changes in the axons of basal forebrain projection neurons.

### BMP4 Restricts Cholinergic Axonal Ramification and Bouton Density

Conditional deletion of *Bmp4* increased VAChT-positive innervation in the neocortex (Fig. 4), with qualitatively denser VAChT immunoreactivity also observed in the hippocampus and amygdala (Fig. 3). Reconstruction of virally labeled cholinergic axons further revealed increased branch points and varicosity density in the neocortex (Fig. 6). Together, these findings indicate that BMP4 restricts axonal ramification and bouton density in neocortical cholinergic projections. In rodents, basal forebrain cholinergic axons form varicosities predominantly in neocortical layers I–III and V, with relatively sparse innervation of layer IV (Ballinger et al., 2016). In *Bmp4* cKO mice, VAChT-positive axons remained sparse within the layer IV barrels, and the VAChT-positive area fraction in layer IV did not differ significantly from that in *Bmp4^fl/fl^* mice (Fig. 4B), although relative VAChT fluorescence intensity in this layer was increased (Fig. 4C). Despite these increases, the characteristic laminar distribution of cholinergic innervation was largely retained.

A previous *in vitro* study demonstrated that BMP4 is secreted near axon terminals and promotes synapse destabilization (Higashi et al., 2018). In that system, BMP4 acted locally at presynaptic sites rather than through a long-range concentration gradient.

Local regulation of axonal organization has also been described for other secreted signaling molecules. Netrin derived from local cellular sources can regulate axon guidance independently of long-range floor-plate-derived cues (Dominici et al., 2017; Varadarajan et al., 2017; Yamauchi et al., 2017), and extracellular Wnt distribution can be spatially restricted through local protein interactions (Takada et al., 2018; Mii et al., 2021). These observations raise the possibility that BMP4 acts within cholinergic projection fields to constrain axonal ramification without substantially altering the overall projection pattern. BMP4 may also act through Smad-dependent transcription, given the enrichment of Smad1/5 phosphorylation in cholinergic neurons (Fig. 2D–F). Future studies examining the spatial range and downstream mechanisms of BMP4 signaling will help determine the relative contributions of local BMP4 action, whether contact-associated or diffusible, and Smad-dependent transcription to cholinergic axonal organization.

Axonal branching and bouton stability are closely linked. Axonal branches often emerge from varicosities, and branch stability depends on the presence and maturation of presynaptic structures (Uesaka et al., 2005; Ruthazer et al., 2006; Yamada et al., 2010). Patterned neuronal activity also contributes to branch stabilization (Matsumoto et al., 2024). The concurrent increases in branch points and varicosity density in *Bmp4* cKO mice are consistent with this link. Given that BMP4 promotes synapse elimination in cultured neurons (Higashi et al., 2018), one possibility is that BMP4 destabilizes nascent presynaptic structures along cholinergic axons and thereby limits the stabilization of axonal branches.

Actin dynamics represent one potential downstream mechanism linking BMP4 signaling to these morphological changes. Both axonal branching and presynaptic stability depend on actin remodeling, which is regulated by pathways involving LIM kinase (LIMK) and actin-depolymerizing factor (ADF)/cofilin (Mizuno, 2013). BMP signaling can regulate growth cone responses through the LIMK–Slingshot–ADF/cofilin pathway (Wen et al., 2007). Examining this pathway in cholinergic axons could test whether altered actin regulation contributes to the increased ramification and varicosity density observed in *Bmp4* cKO mice.

Glial cells may also contribute to the regulation of cholinergic axonal structure. Microglia and astrocytes eliminate dysfunctional or excess synapses (Wilton et al., 2019; Kurematsu et al., 2022; Morizawa et al., 2022) and could therefore influence axonal morphology indirectly through synaptic remodeling. BMP4 suppresses synapse formation even in neuron-enriched cultures in which glial proliferation is strongly suppressed by cytosine arabinoside (Higashi et al., 2018), supporting the existence of a direct neuronal mechanism, at least *in vitro*. Nevertheless, interactions between neurons and glia may further shape cholinergic axonal organization *in vivo*.

### Layer-dependent Effects of BMP4 on Cholinergic Axons

In *Bmp4* cKO mice, axonal branch points showed a significant genotype × layer interaction, with a significant increase detected in layer V (Fig. 6E), whereas varicosity density increased overall with no significant genotype × layer interaction (Fig. 6F).

Layer V contains the principal corticofugal projection neurons (Harris and Shepherd, 2015). Increased cholinergic ramification in this layer may particularly influence cortical output. The laminar organization of the neocortex follows a common circuit pattern that is modified in an area-specific manner (Harris and Shepherd, 2015). As axons from the frontal–parietal, parietal–occipital, and temporal areas were included in the present analysis (Fig. 6C), whole-brain mapping of individual cholinergic axons will be useful for determining whether the layer-dependent effects observed here vary across neocortical areas.

### Functional Implications

Cholinergic projections from the basal forebrain modulate cortical plasticity and cognitive processes (Wenk, 1997; Hasselmo, 1999; Xu et al., 2015; Guo et al., 2019). The increased axonal branching and bouton density in *Bmp4* cKO mice may increase the number of potential acetylcholine release sites in the neocortex and thereby influence cortical activity and cognitive performance. Brain-wide activity imaging and behavioral analyses will be required to determine the functional consequences of these structural changes in cholinergic projections. Cortical circuit function also depends on E/I balance, which can be regulated by BMP–Smad signaling (Okur et al., 2024). In *Bmp4* cKO mice, the area fractions of VGluT1- and VGAT-positive signals in neocortical layer II/III did not differ significantly from those in *Bmp4^fl/fl^* mice (Fig. 2A–C). Electron microscopy or super-resolution imaging could reveal whether finer changes in synaptic organization accompany the increased cholinergic innervation. In addition, the adult experiments in this study were performed in males; whether BMP4 similarly regulates cholinergic axonal ramification in females remains to be determined.

In conclusion, BMP4 expressed in cholinergic neurons restricts the axonal ramification and bouton density of basal forebrain cholinergic projections. The enrichment of BMP– Smad signaling in these neurons during early postnatal development suggests that BMP4 constrains cholinergic innervation during postnatal circuit maturation.

## Author Contributions

T.H., Y.K., and K.N. designed research; T.H. performed research; A.S. and S.O. contributed unpublished reagents/analytic tools; T.H., Y.K., and K.N. analyzed data; T.H., Y.K., K.N., and S.O. wrote the paper.

## Acknowledgments

This work was supported by JSPS KAKENHI (Grant Number JP19K16282 to T.H.) and a National Defense Medical College Research Grant. The ChAT-Cre mouse line (STOCK Tg(Chat-cre)GM24Gsat/Mmucd; RRID:MMRRC_017269-UCD) was obtained from the Mutant Mouse Resource and Research Center (MMRRC) at the University of California, Davis, an NIH-funded strain repository, and was donated to the MMRRC by Nathaniel Heintz, Ph.D., The Rockefeller University, GENSAT, and Charles Gerfen, Ph.D., National Institutes of Health, National Institute of Mental Health. We thank Dr. Hongkui Zeng for depositing the pCAG-FLEX-EGFP-WPRE plasmid with Addgene (viral prep #51502-AAV9; RRID:Addgene_51502). We also thank Mayumi Watanabe for technical assistance with tissue preparation and histological analyses, and Natsumi Masumoto for preparing the visual abstract. The authors used ChatGPT (OpenAI) and Claude (Anthropic) during manuscript preparation to assist with language refinement. The manuscript was subsequently reviewed and edited by Editage (www.editage.com). All scientific content, interpretations, and conclusions were reviewed and approved by the authors.

